## supplemental figures for "A microbial metabolite protects against graft-versus-host disease via mTORC1 and STING-dependent intestinal regeneration"

**Table S1**

| | | |
|---|---|---|
| **Patients – no.** | | 50 |
| **Mean age at HSCT – yrs ± SE** | | 56.5 ± 1.5 |
| **Male/Female Sex – no. (%)** | | 27 (54)/23 (46) |
| **Diagnosis – no. (%)** | Acute leukaemia | 22 (44) |
| | MDS/MPN | 19 (38) |
| | NHL | 5 (10) |
| | other | 4 (8) |
| **Donor type – no. (%)** | Unrelated | 35 (70) |
| | Sibling | 5 (10) |
| | Haploidentical | 10 (20) |
| **Conditioning – no. (%)** | Ablative | 7 (14) |
| | Reduced intensity | 43 (86) |
| **Stem cell source – no. (%)** | PBSC | 42 (84) |
| | BM | 8 (16) |
| **High DAT levels – no. (%)** | | 22 (44) |
| **High ICA levels – no. (%)** | | 10 (20) |
| **2 year relapse incidence (%)** | | 10 (20) |
| **2 year TRM incidence (%)** | | 10 (20) |
| **2 year survival (%)** | | 31 (62) |

**Patient characteristics.** HSCT denotes hematopoietic stem cell transplantation, SE standard error, MDS myelodysplastic syndrome, MPN myeloproliferative neoplasia, NHL Non-Hodgkin lymphoma, PBSC peripheral-blood stem cell, BM bone marrow, and TRM transplantation-related mortality. Diseases categorized as “other” include aplastic anemia and myelosarcoma.

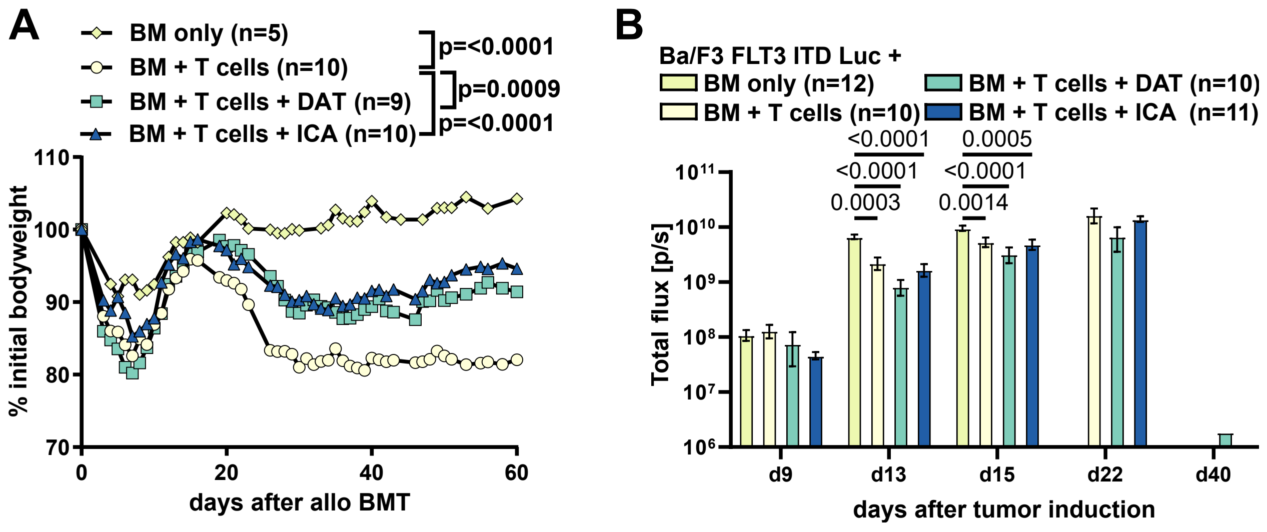

A
BM only (n=5)
BM + T cells (n=10)
BM + T cells + DAT (n=9)
BM + T cells + ICA (n=10)
p=<0.0001
p=0.0009
p=<0.0001
% initial bodyweight
110
100
90
80
70
0
20
40
60
days after allo BMT
B
Ba/F3 FLT3 ITD Luc +
BM only (n=12)
BM + T cells (n=10)
BM + T cells + DAT (n=10)
BM + T cells + ICA (n=11)
<0.0001
<0.0001
0.0003
0.0005
<0.0001
0.0014
Total flux [p/s]
10^11
10^10
10^9
10^8
10^7
10^6
d9
d13
d15
d22
d40
days after tumor induction


### Figure S1

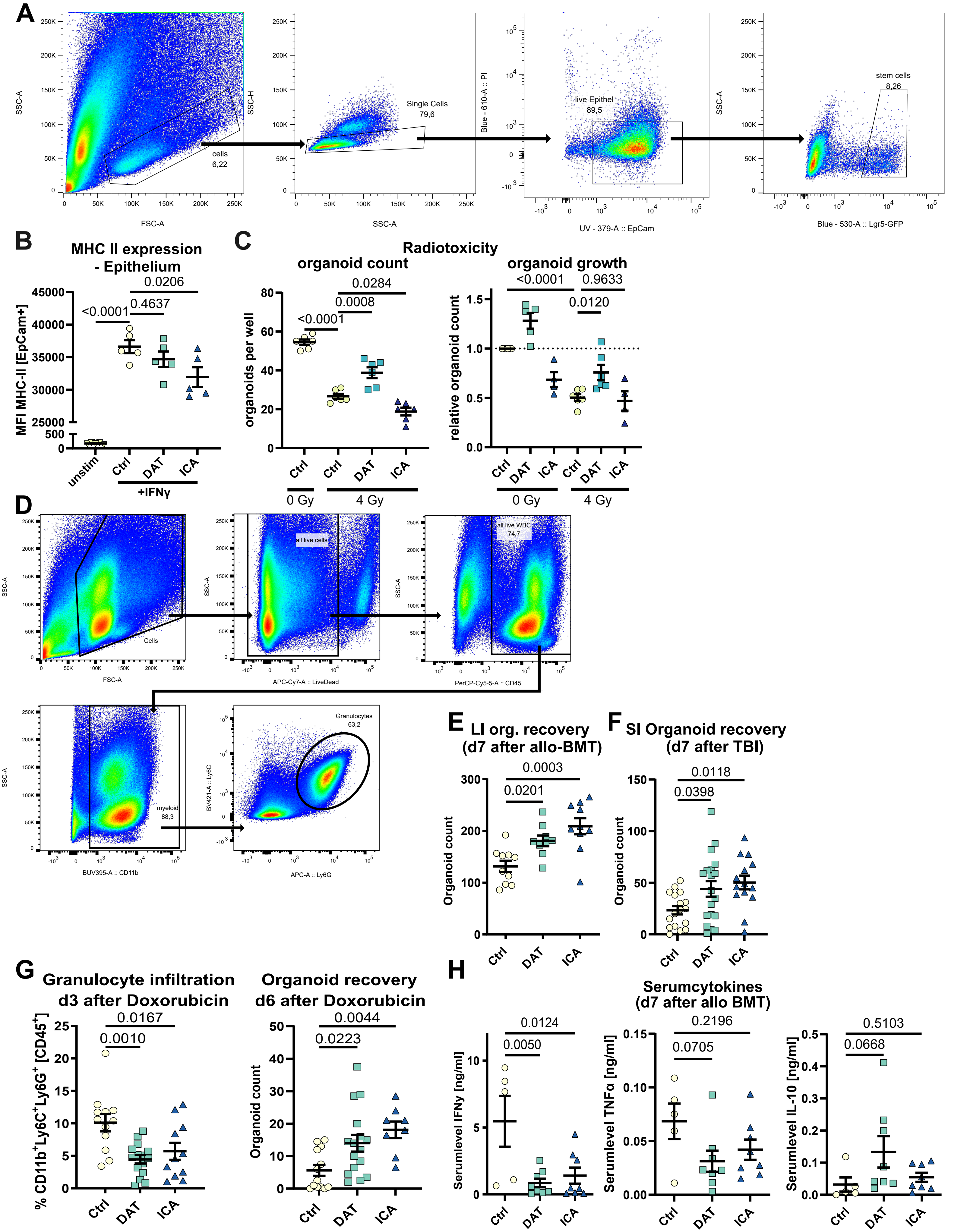
A
SSC-A
FSC-A
cells 6,22
SSC-H
Single Cells 79,6
Blue - 610-A :: PI
live Epithel 89,5
UV - 379-A :: EpCam
stem cells 8,26
Blue - 530-A :: Lgr5-GFP
B
MHC II expression - Epithelium
MFI MHC-II [EpCam+]
<0.0001
0.4637
0.0206
unstim
Ctrl
DAT
ICA
+IFNγ
C
Radiotoxicity
organoid count
organoids per well
<0.0001
0.0008
0.0284
0 Gy
4 Gy
organoid growth
relative organoid count
<0.0001
0.9633
0.0120
D
Cells
all live cells
APC-Cy7-A :: LiveDead
all live WBC 74,7
PerCP-Cy5-5-A :: CD45
myeloid 88,3
BUV395-A :: CD11b
Granulocytes 63,2
BV421-A :: Ly6C
APC-A :: Ly6G
E
LI org. recovery (d7 after allo-BMT)
Organoid count
0.0201
0.0003
F
SI Organoid recovery (d7 after TBI)
0.0398
0.0118
G
Granulocyte infiltration d3 after Doxorubicin
% CD11b+Ly6C+Ly6G+ [CD45+]
0.0010
0.0167
Organoid recovery d6 after Doxorubicin
0.0223
0.0044
H
Serumcytokines (d7 after allo BMT)
Serumlevel IFNγ [ng/ml]
0.0050
0.0124
Serumlevel TNFα [ng/ml]
0.0705
0.2196
Serumlevel IL-10 [ng/ml]
0.0668
0.5103

**Figure S2**

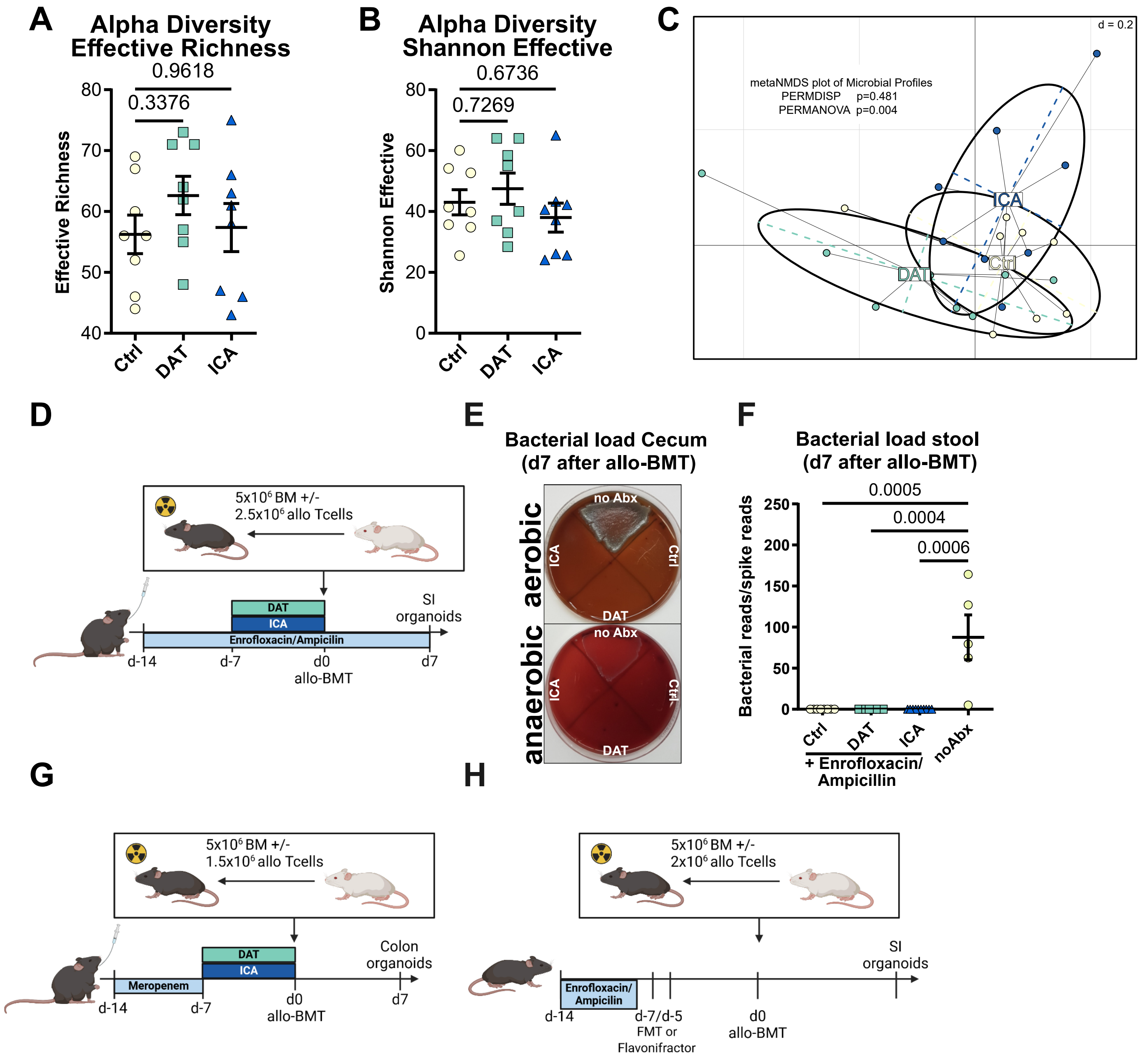
A
Alpha Diversity
Effective Richness
0.9618
0.3376
Effective Richness
Ctrl
DAT
ICA
B
Alpha Diversity
Shannon Effective
0.6736
0.7269
Shannon Effective
C
d = 0.2
metaNMDS plot of Microbial Profiles
PERMDISP p=0.481
PERMANOVA p=0.004
ICA
DAT
Ctrl
D
5x10⁶ BM +/-
2.5x10⁶ allo Tcells
DAT
ICA
Enrofloxacin/Ampicilin
SI organoids
d-14
d-7
d0
allo-BMT
d7
E
Bacterial load Cecum
(d7 after allo-BMT)
aerobic
anaerobic
no Abx
ICA
Ctrl
DAT
F
Bacterial load stool
(d7 after allo-BMT)
Bacterial reads/spike reads
0.0005
0.0004
0.0006
Ctrl
DAT
ICA
noAbx
+ Enrofloxacin/
Ampicillin
G
5x10⁶ BM +/-
1.5x10⁶ allo Tcells
Meropenem
DAT
ICA
Colon organoids
d-14
d-7
d0
allo-BMT
d7
H
5x10⁶ BM +/-
2x10⁶ allo Tcells
Enrofloxacin/
Ampicilin
SI organoids
d-14
d-7/d-5
FMT or
Flavonifractor
d0
allo-BMT

### Figure S3

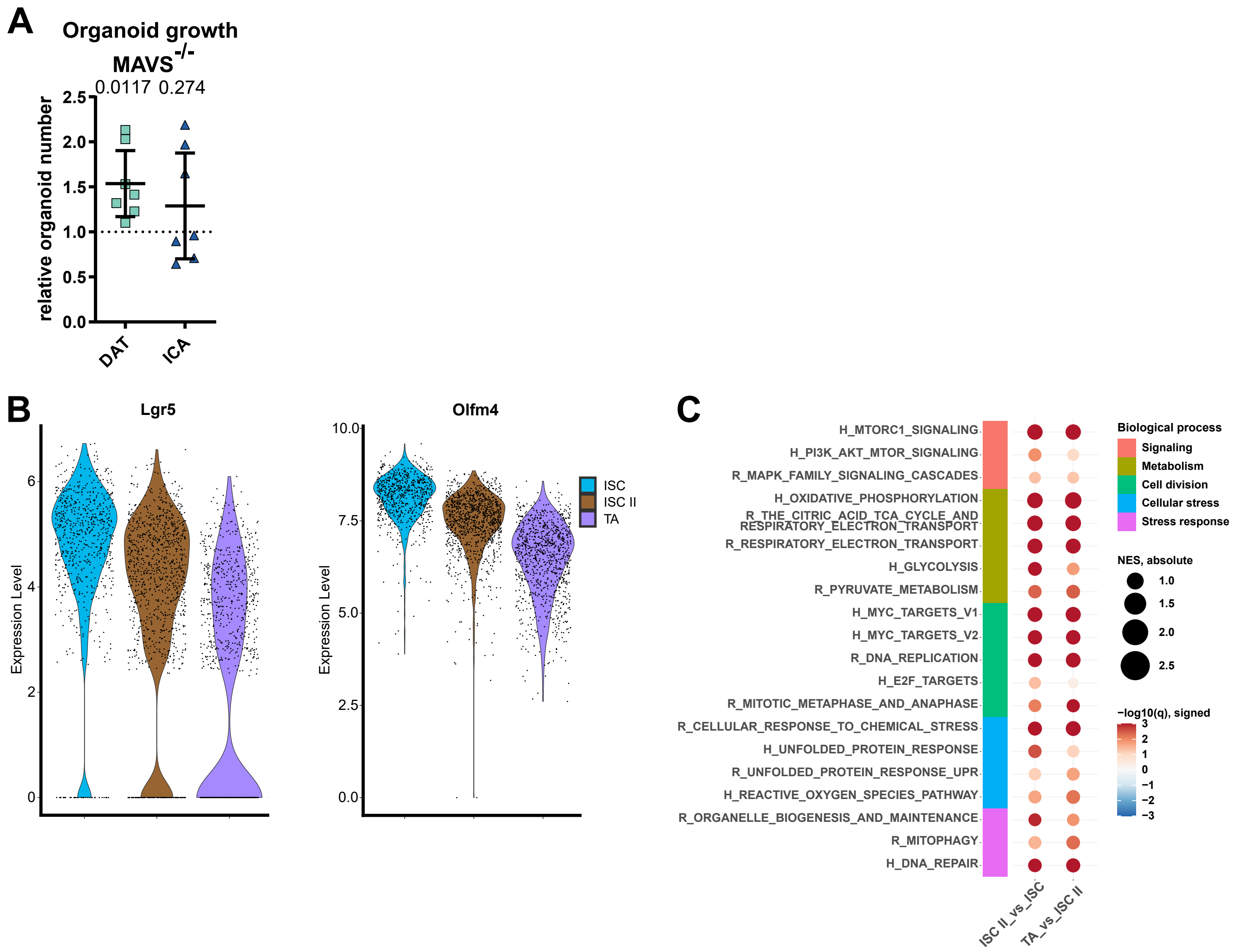
A
Organoid growth
MAVS-/-
0.0117 0.274
relative organoid number
2.5
2.0
1.5
1.0
0.5
0.0
DAT
ICA
B
Lgr5
Olfm4
Expression Level
ISC
ISC II
TA
C
H_MTORC1_SIGNALING
H_PI3K_AKT_MTOR_SIGNALING
R_MAPK_FAMILY_SIGNALING_CASCADES
H_OXIDATIVE_PHOSPHORYLATION
R_THE_CITRIC_ACID_TCA_CYCLE_AND RESPIRATORY_ELECTRON_TRANSPORT
R_RESPIRATORY_ELECTRON_TRANSPORT
H_GLYCOLYSIS
R_PYRUVATE_METABOLISM
H_MYC_TARGETS_V1
H_MYC_TARGETS_V2
R_DNA_REPLICATION
H_E2F_TARGETS
R_MITOTIC_METAPHASE_AND_ANAPHASE
R_CELLULAR_RESPONSE_TO_CHEMICAL_STRESS
H_UNFOLDED_PROTEIN_RESPONSE
R_UNFOLDED_PROTEIN_RESPONSE_UPR
H_REACTIVE_OXYGEN_SPECIES_PATHWAY
R_ORGANELLE_BIOGENESIS_AND_MAINTENANCE
R_MITOPHAGY
H_DNA_REPAIR
ISC II_vs_ISC
TA_vs_ISC II
Biological process
Signaling
Metabolism
Cell division
Cellular stress
Stress response
NES, absolute
1.0
1.5
2.0
2.5
−log10(q), signed
3
2
1
0
−1
−2
−3

### Figure S4

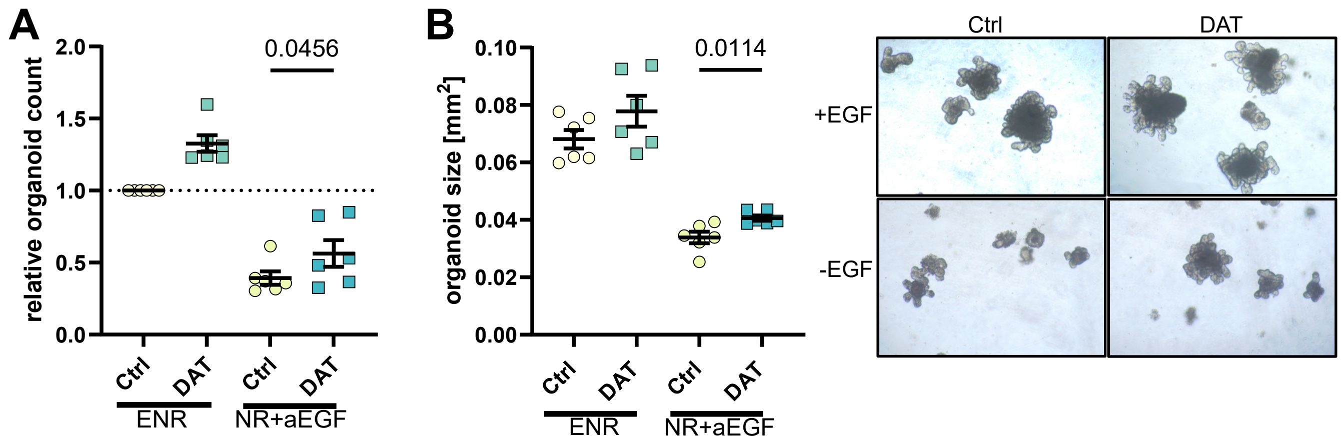
A
relative organoid count
2.0
1.5
1.0
0.5
0.0
0.0456
Ctrl
DAT
Ctrl
DAT
ENR
NR+aEGF
B
organoid size [mm2]
0.10
0.08
0.06
0.04
0.02
0.00
0.0114
Ctrl
DAT
Ctrl
DAT
ENR
NR+aEGF
Ctrl
DAT
+EGF
-EGF

Figure S5

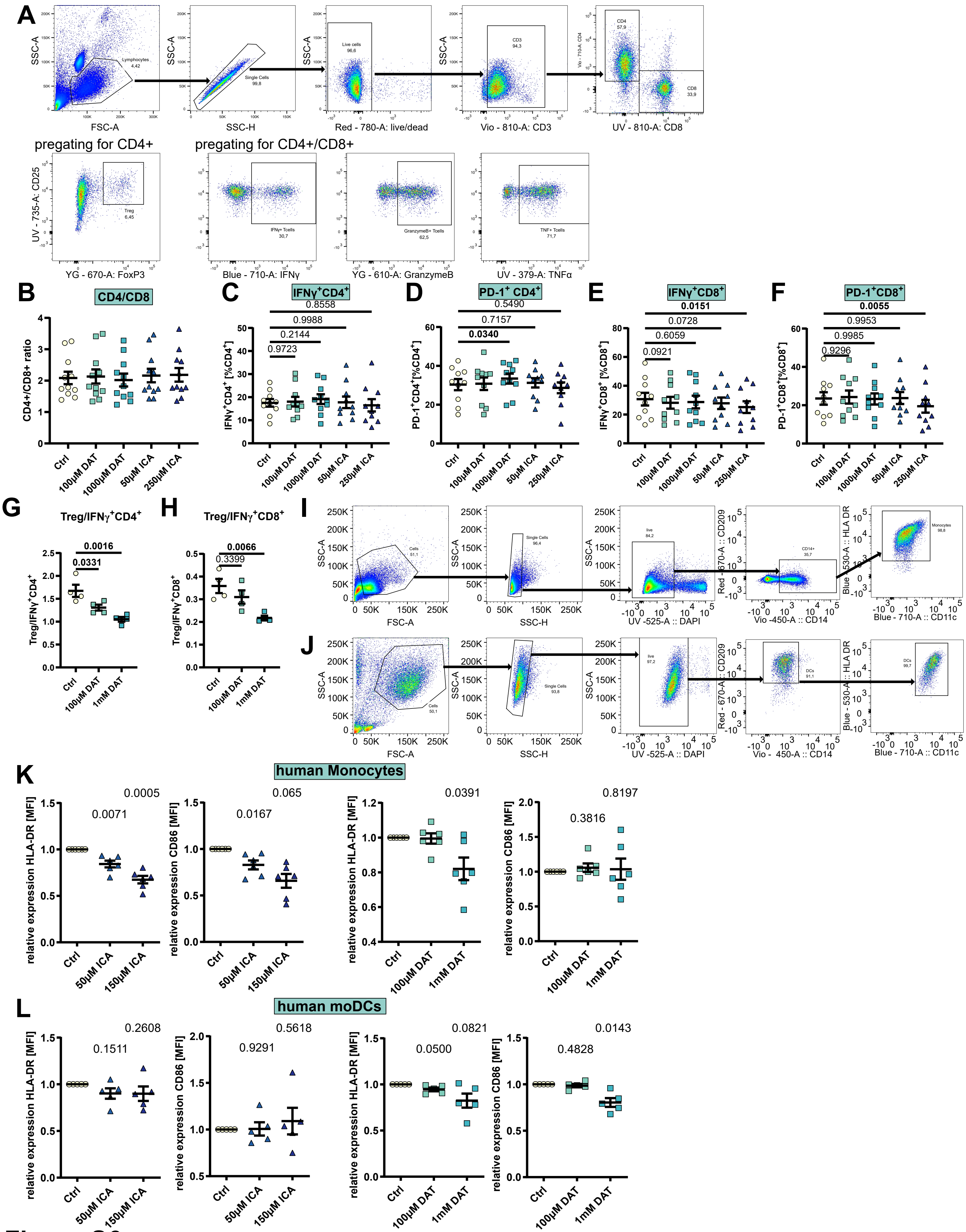
A
pregating for CD4+
pregating for CD4+/CD8+
B
CD4/CD8
C
IFNγ+CD4+
D
PD-1+ CD4+
E
IFNγ+CD8+
F
PD-1+CD8+
G
Treg/IFNγ+CD4+
H
Treg/IFNγ+CD8+
I
J
K
human Monocytes
L
human moDCs

Figure S6

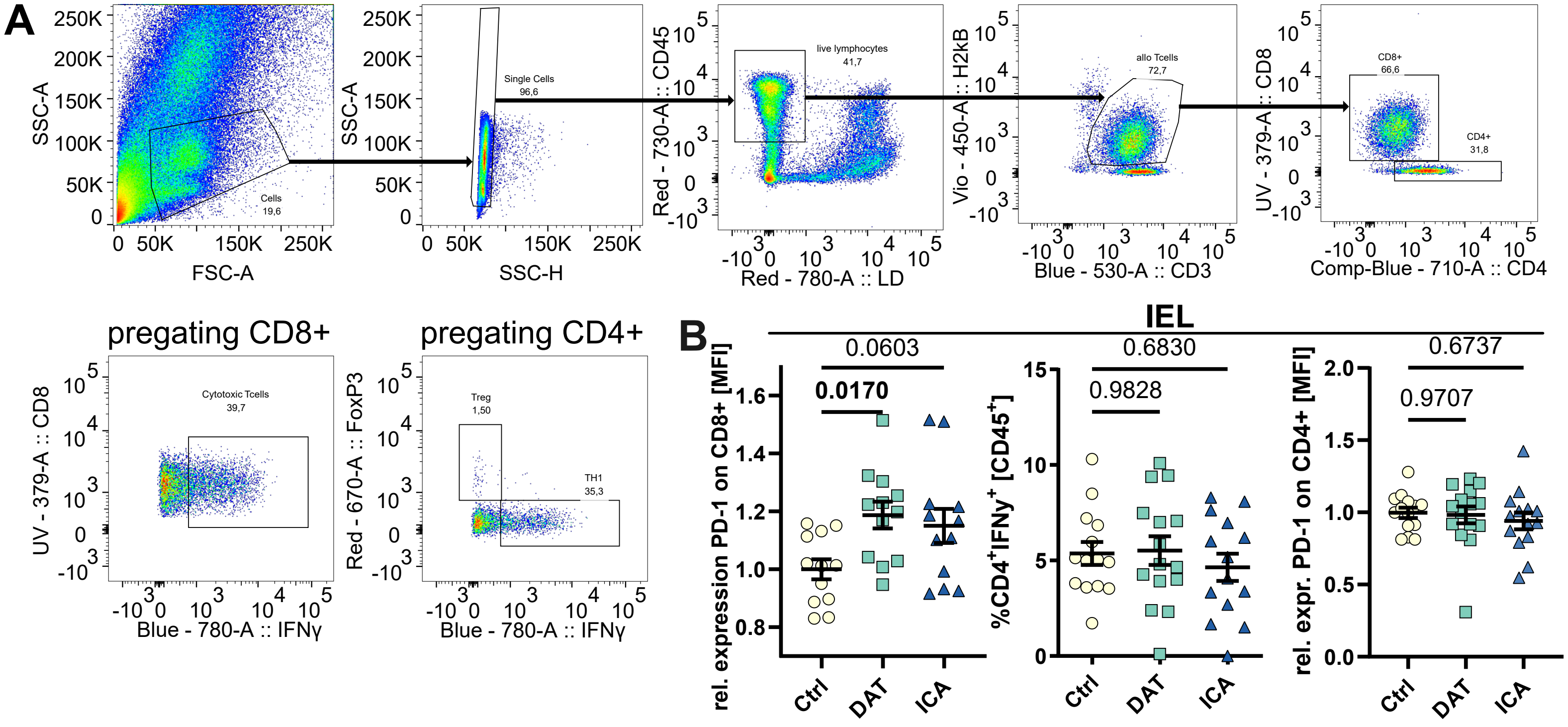
A
SSC-A
FSC-A
Cells
19,6
SSC-H
Single Cells
96,6
Red - 730-A :: CD45
Red - 780-A :: LD
live lymphocytes
41,7
Vio - 450-A :: H2kB
Blue - 530-A :: CD3
allo Tcells
72,7
UV - 379-A :: CD8
Comp-Blue - 710-A :: CD4
CD8+
66,6
CD4+
31,8
pregating CD8+
Cytotoxic Tcells
39,7
Blue - 780-A :: IFNγ
pregating CD4+
Red - 670-A :: FoxP3
Treg
1,50
TH1
35,3
B
IEL
rel. expression PD-1 on CD8+ [MFI]
0.0603
0.0170
%CD4+IFNy+ [CD45+]
0.6830
0.9828
rel. expr. PD-1 on CD4+ [MFI]
0.6737
0.9707
Ctrl
DAT
ICA

**Figure S7**

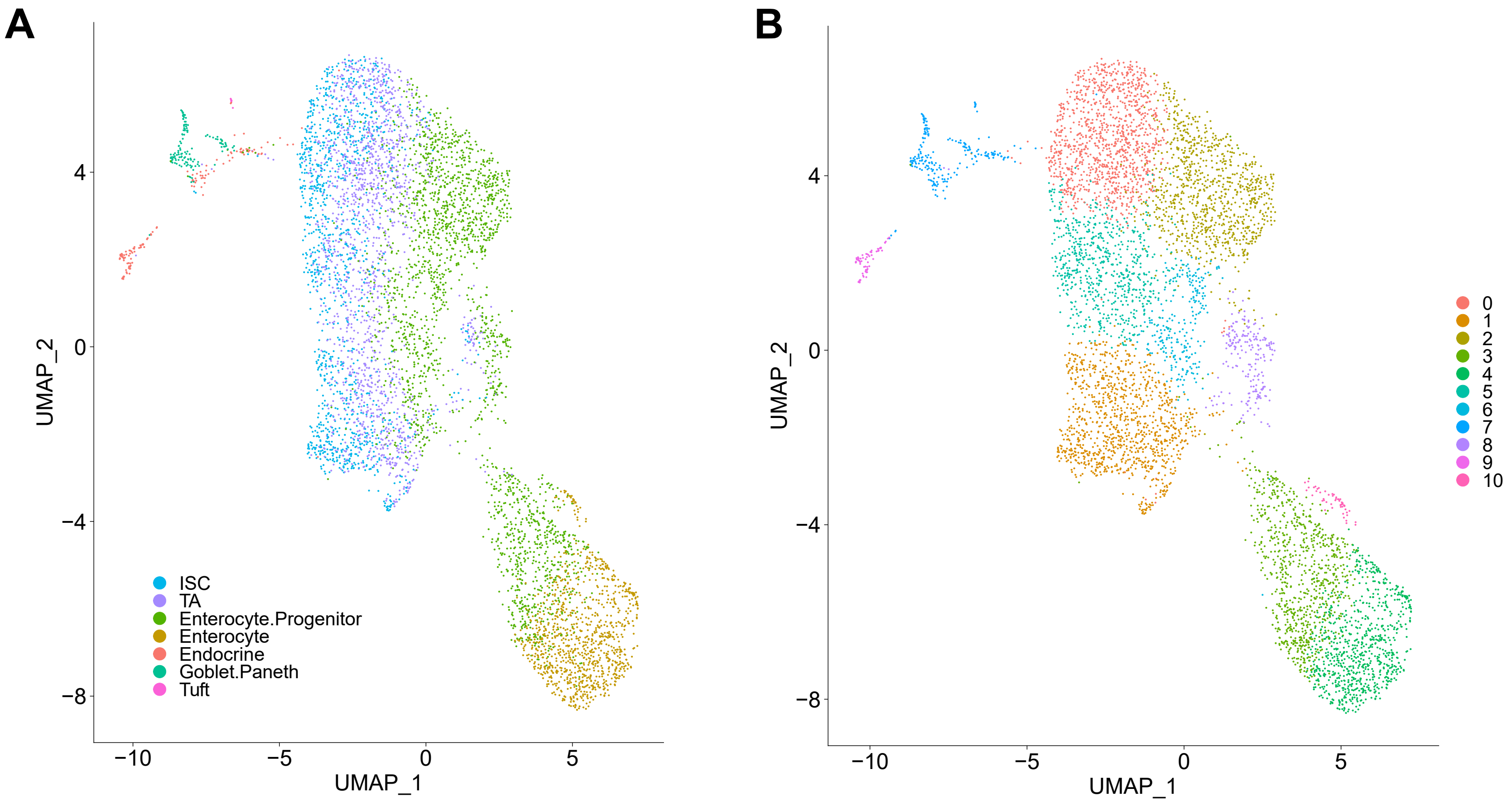


**Figure S8, related to methods**

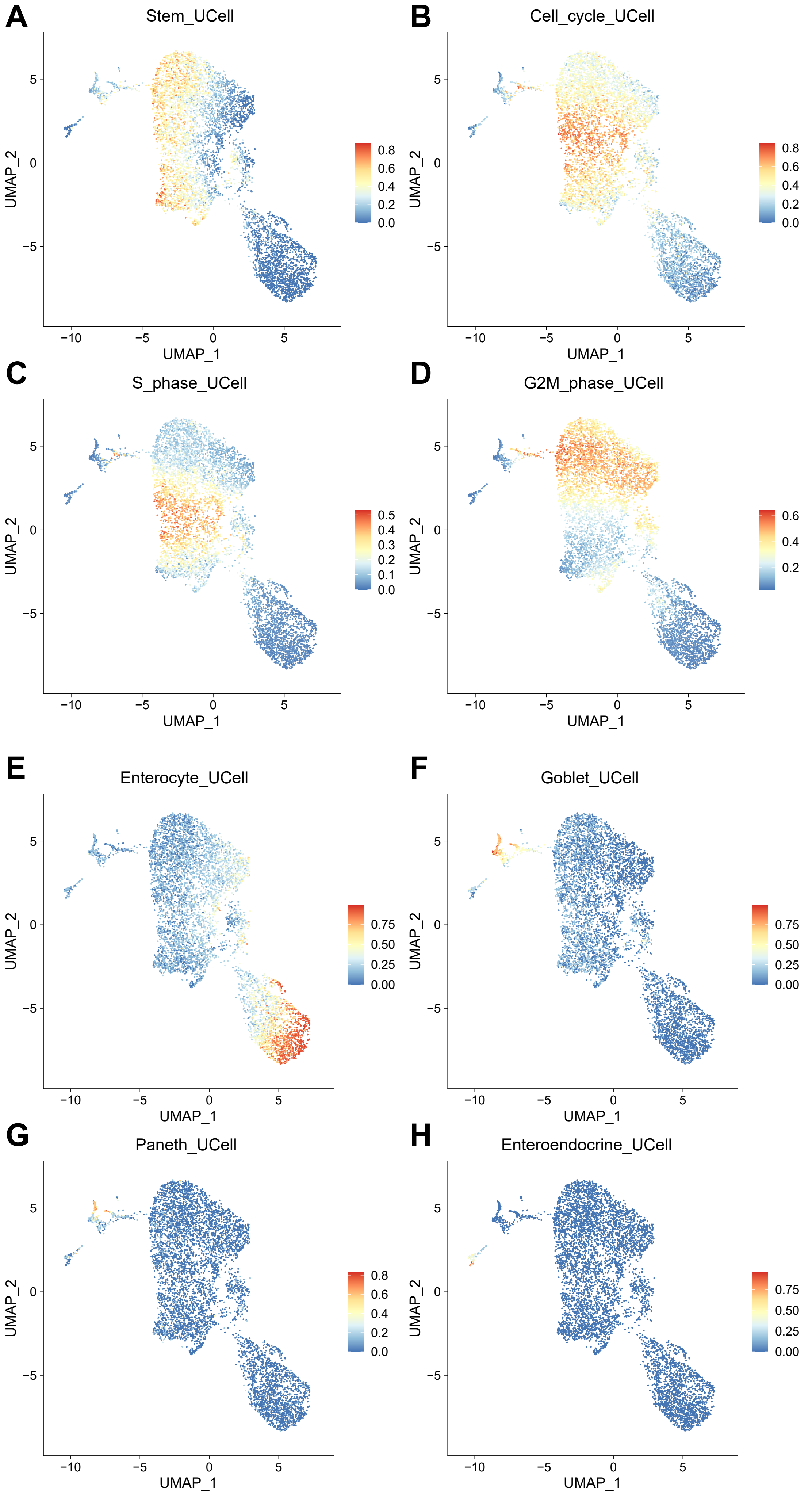


**Figure S9, related to methods**

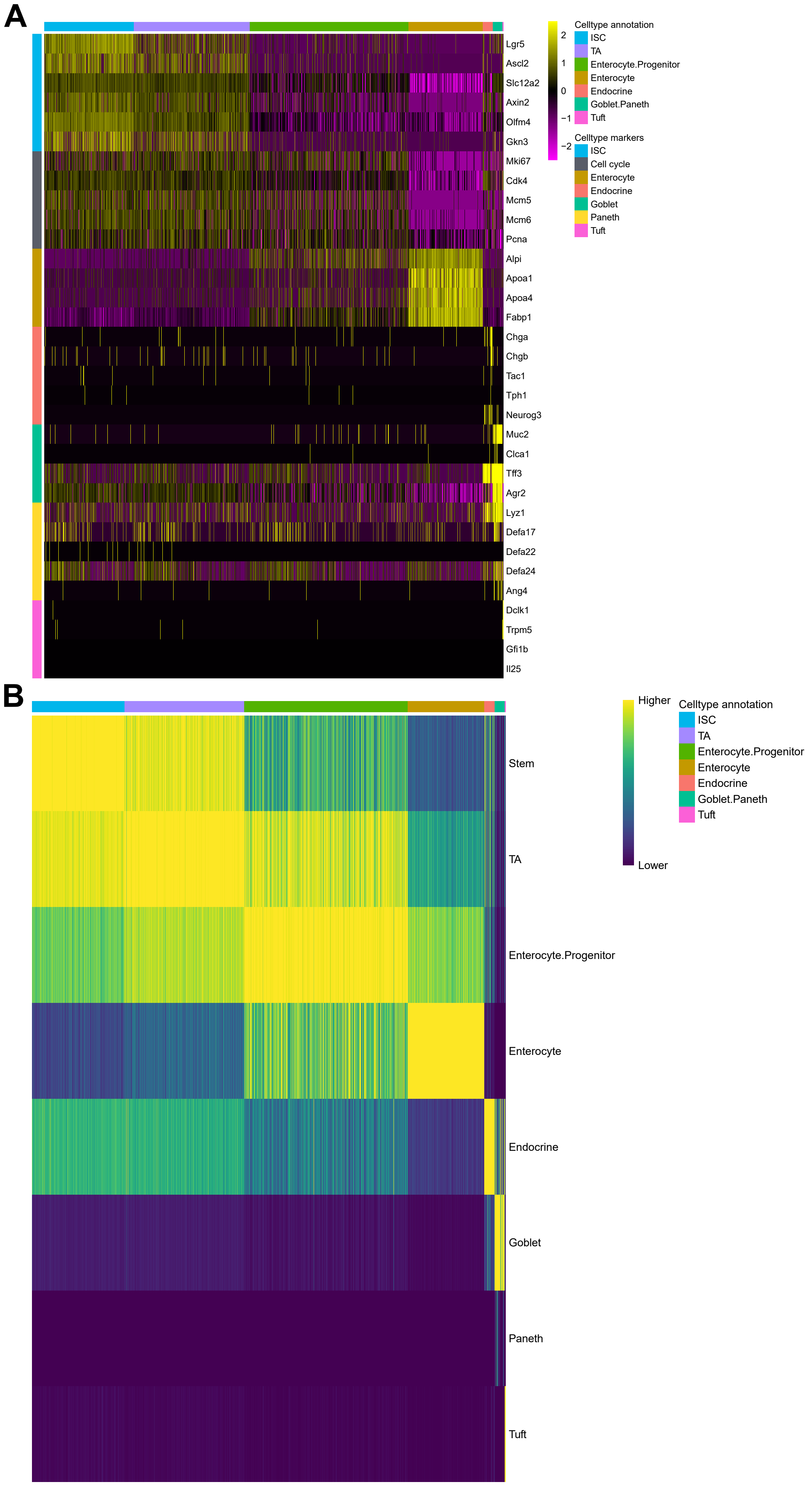


**Figure S10, related to methods**

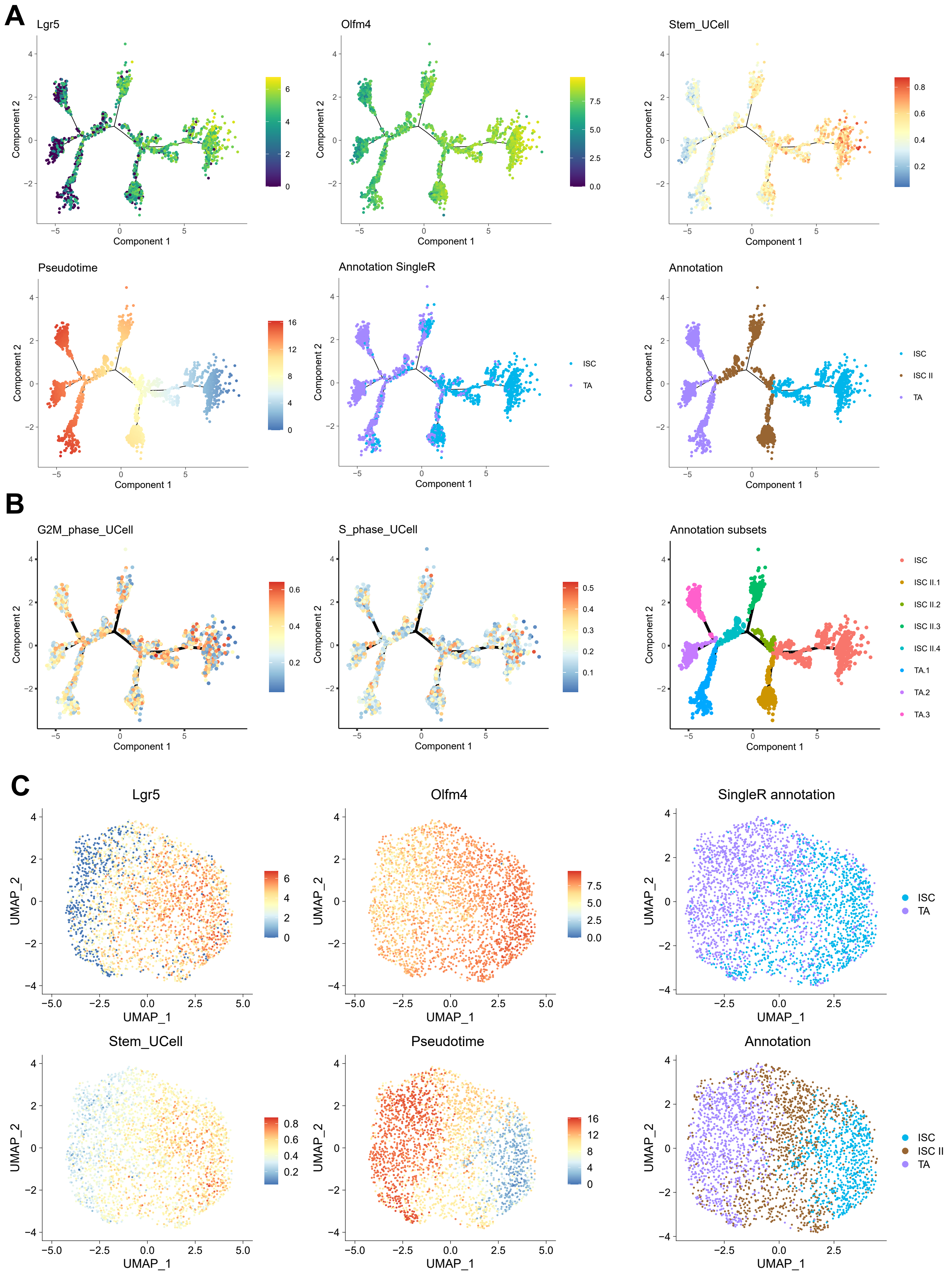


### Figure S11, related to methods

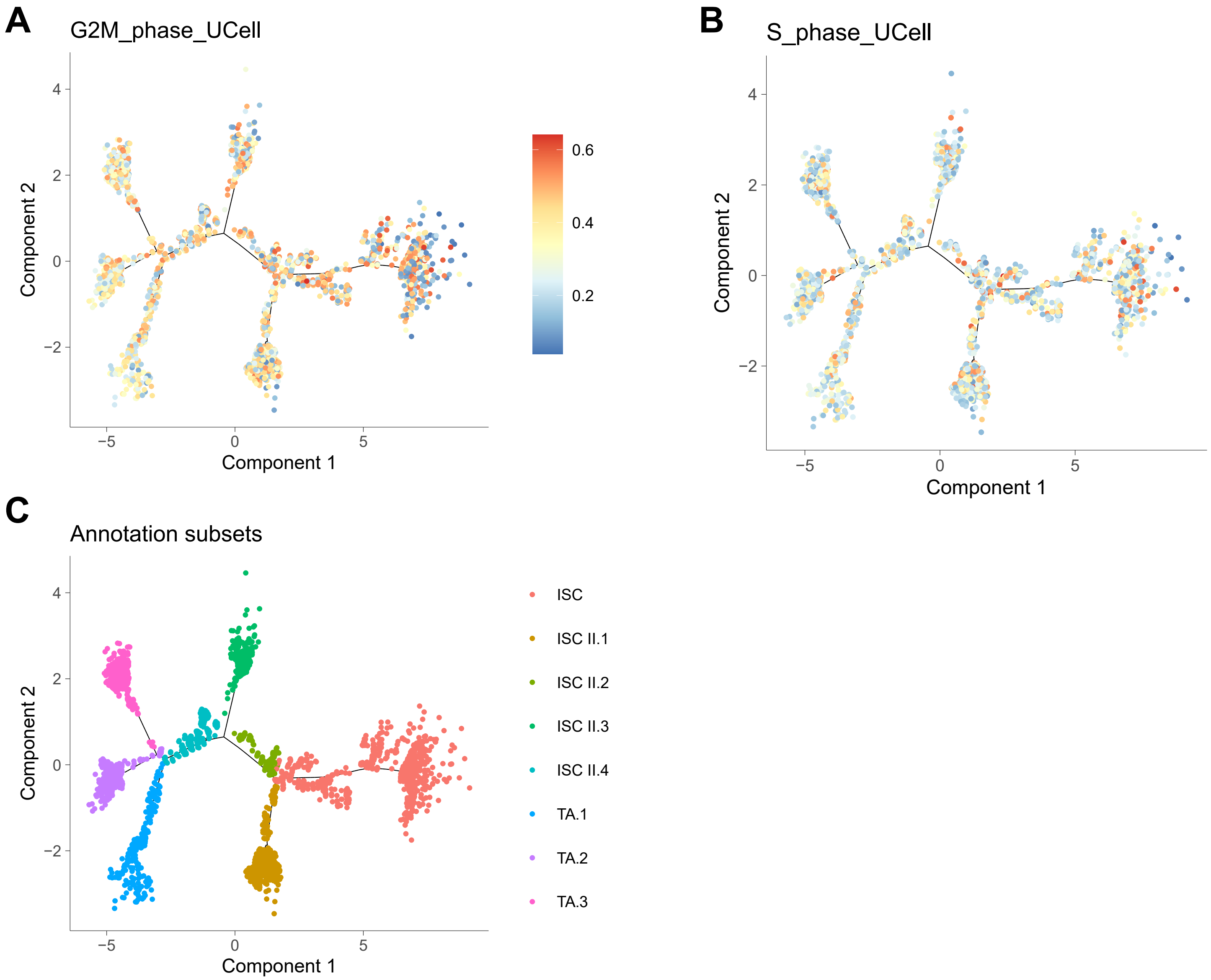
A
G2M_phase_UCell
Component 2
Component 1
0.6
0.4
0.2
B
S_phase_UCell
Component 2
Component 1
0.5
0.4
0.3
0.2
0.1
C
Annotation subsets
Component 2
Component 1
ISC
ISC II.1
ISC II.2
ISC II.3
ISC II.4
TA.1
TA.2
TA.3

Figure S12
